# Loss of cerebellar function selectively affects intrinsic rhythmicity of eupneic breathing

**DOI:** 10.1101/784769

**Authors:** Yu Liu, Shuhua Qi, Fridtjof Thomas, Brittany L. Correia, Angela P. Taylor, Roy V. Sillitoe, Detlef H. Heck

## Abstract

Respiration is controlled by central pattern generating circuits in the brain stem, whose activity can be modulated by inputs from other brain areas to adapt respiration to autonomic and behavioral demands. The cerebellum is known to be part of the neuronal circuitry activated during respiratory challenges, such as hunger for air, but has not been found to be involved in the control of unobstructed breathing at rest (eupnea). Here we applied a measure of intrinsic rhythmicity, the CV2, which evaluates the similarity of subsequent intervals and is thus sensitive to changes in rhythmicity at the temporal resolution of individual respiratory intervals. The variability of intrinsic respiratory rhythmicity was reduced in a mouse model of cerebellar ataxia compared to their healthy littermates. Irrespective of that difference, the average respiratory rate and the average coefficient of variation (CV) were comparable between healthy and ataxic mice. We argue that these findings are consistent with a proposed role of the cerebellum in the coordination of respiration with other rhythmic orofacial movements, such as fluid licking and swallowing.

## INTRODUCTION

The cerebellum has extensive reciprocal connections with the brain stem and cerebellar neuropathology is known to affect brain stem controlled processes, such as cardiovascular and respiratory function (Harper et al., 1998; Paton et al., 1991; Rector et al., 2006). The involvement of the cerebellum in respiration seems to be central to respiratory challenges, such as hypoxia or hypercapnia (Macey et al., 2005; Parsons et al., 2001). Neurons in the medial cerebellar nuclei project to brain stem areas containing the respiratory pattern generating circuits (Lu et al., 2013) and neuronal activity in the medial cerebellar nuclei is entrained by the respiratory rhythm (Lu et al., 2013; Xu and Frazier, 2002). Investigations into eupneic respiration, however, failed to implicate the cerebellum in animal models (Moruzzi, 1940; Xu and Frazier, 2002; Xu et al., 1995) or humans (Ebert et al., 1995). Previously published findings suggested that ablation of the fastigial nucleus altered respiratory responses to hypercapnia but did not alter eupneic breathing (Xu and Frazier, 2002). However, a recent study conducted in juvenile mice reported the possibility of only a modest influence of cerebellar cortical function during respiration as determined by a measure of breathing regularity called inter breath interval (van der Heijden and Zoghbi, 2018). Collectively, these previous studies that investigated the involvement of the cerebellum in respiration have been complicated by the use of anaesthetized conditions or analyses conducted before postnatal day 30, an age prior to which functional cerebellar circuits have yet to reach maturity (Arancillo et al., 2015). We therefore sought to test the role of the mature cerebellum in eupneic breathing in awake conditions using multiple physiological measures of system rhythmicity.

To quantitatively address this problem, we compared the average respiratory rate, coefficient of variation (CV) of the respiratory rhythm and the intrinsic rhythmicity of respiration (CV2) (Holt et al., 1996) of eupneic breathing in healthy mice (controls, CT) and in adult mice with cerebellar ataxia. Cerebellar ataxia was induced using the Cre/LoxP genetic approach to selectively block Purkinje cell GABAergic neurotransmission (*L7^Cre^;Vgat^flox/flox^*), thereby functionally disconnecting the cerebellar cortex from the cerebellar nuclei (White et al., 2014). Spontaneous respiratory behavior was measured in a plethysmograph.

Cerebellar ataxia did not affect the average respiratory rate or the CV of the respiratory rhythm. However, compared to their CT littermates, *L7^Cre^;Vgat^flox/flox^* mutant mice (MU) showed increased intrinsic rhythmicity, as measured by the CV2. The CV2 evaluates the similarity of pairs of intervals, and is thus sensitive to brief changes in rhythmicity at a temporal resolution of individual interval durations. The CV2 provides a measure of intrinsic rhythmicity in the sense that this measure is sensitive to short periods of highly regular breathing but less sensitive to slow rate variations than the CV. Based on our results and findings from previously published studies, we propose a role for the cerebellum in the coordination of multiple rhythmic orofacial movements, such as coordinating respiration with swallowing.

## RESULTS

To measure spontaneous respiratory behavior, mice were placed in a plethysmograph chamber, where they could move freely (Fig. 1). Spontaneous respiration was monitored for 30 min and compared between MU mice and their CT litter mates (Table 1). Male and female mice of both phenotypes were divided into two age groups, 2-3 or 5-7 months old (Table 1), corresponding approximately to the transition from adolescence/young adults to mature adults in human development (Flurkey et al., 2007).

**Figure 1:**
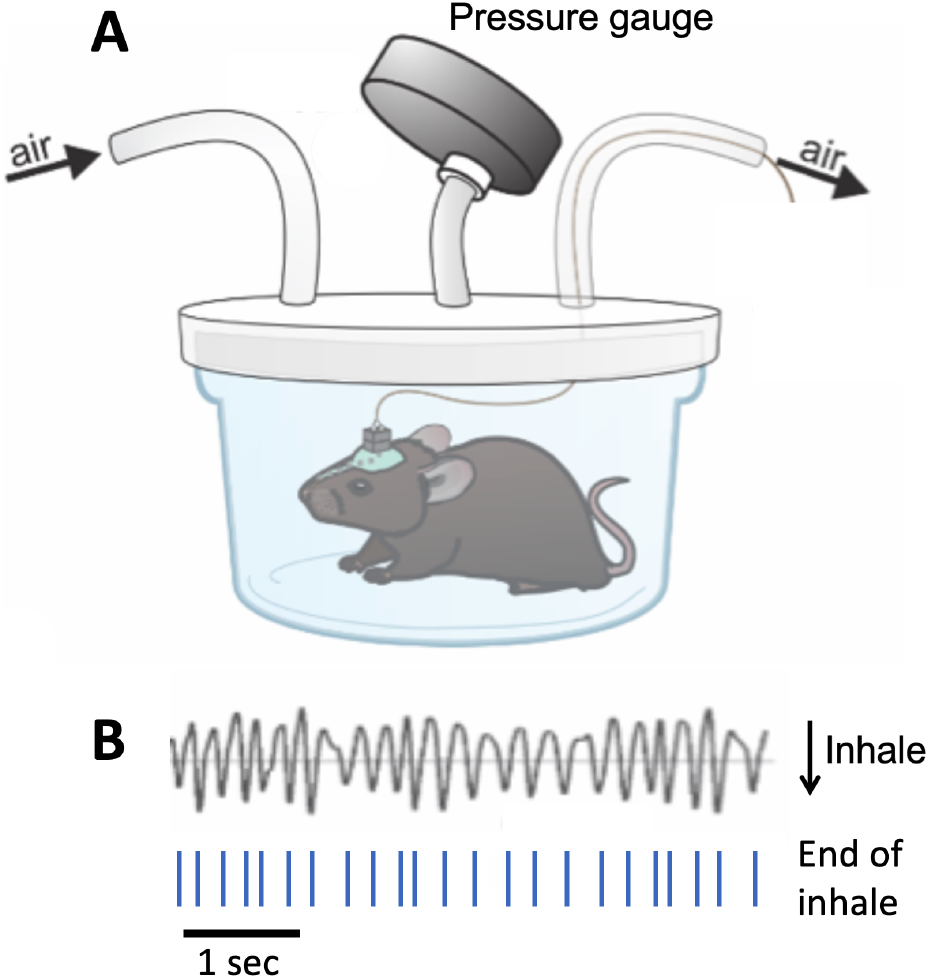
Mouse respiratory behavior was measured using a custom-made plethysmograph chamber. **A)** Pressure inside the chamber was measured with a pressure transducer (Validyne Engineering, USA). Inhalation movements increase the pressure inside the chamber, resulting in decreases in voltage. **B)** Raw voltage output from the pressure transducer reflecting respiration related pressure changes. Inhalation related increases in pressure are reflected as decreases in voltage. Troughs in the voltage signal reflect the end of inhale times, which were marked and used as temporal aligns for further analysis of the respiratory rhythm.

**Table 1:**
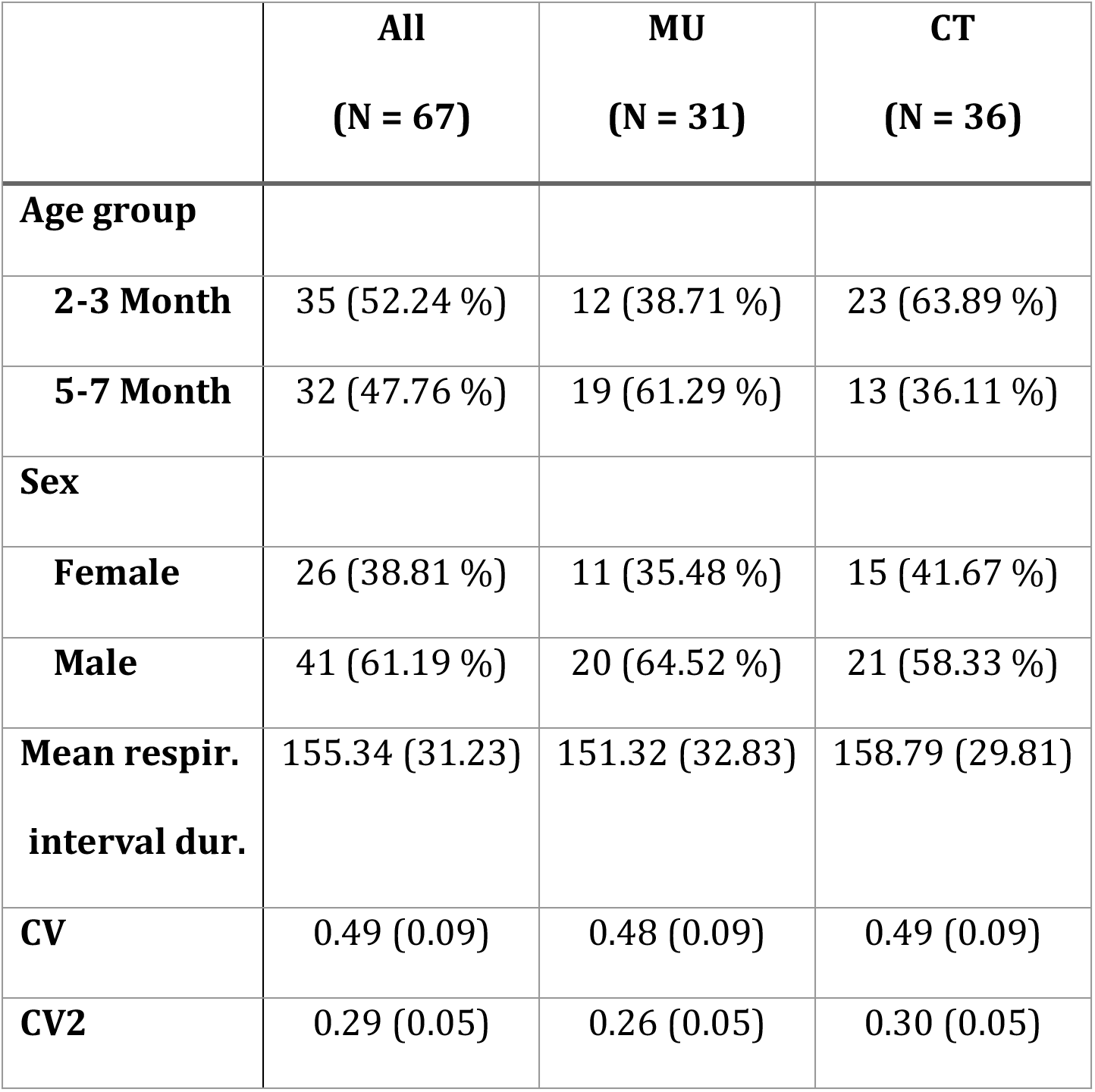
Mouse group compositions and phenotypes analyzed. Shown are count and (%) or mean and (std.dev.), respectively, depending on whether the variable is discrete or continuous.

Respiratory activity in the plethysmograph chamber causes rhythmic pressure changes, which were measured using a pressure transducer (Fig. 1A). Rhythmic decreases in voltage output from the transducer corresponded to inhalation movements, and the times of voltage trough minima were marked as “end-of-inhalation” times (Fig. 1B). All further analysis was based on these temporal markers.

The average respiratory rate was evaluated by calculating the average duration of inter-inhalation intervals. Comparison of mean interval durations between MU mice and CT littermates within age groups revealed no difference between groups (p=0.99, adjusted for sex and age; Fig. 2 - see Materials and Methods/Statistical analysis for details). Older mice, however, had shorter average interval durations than the younger mice (p<0.001; adjusted for sex), consistent with a known age-related increase in respiratory rate (Rodriguez-Molinero et al., 2013). Comparison of respiratory rate by sex showed a systematic, albeit not very pronounced difference (p=0.048, adjusted for age), with female mice having higher respiratory rates than males.

**Figure 2).**
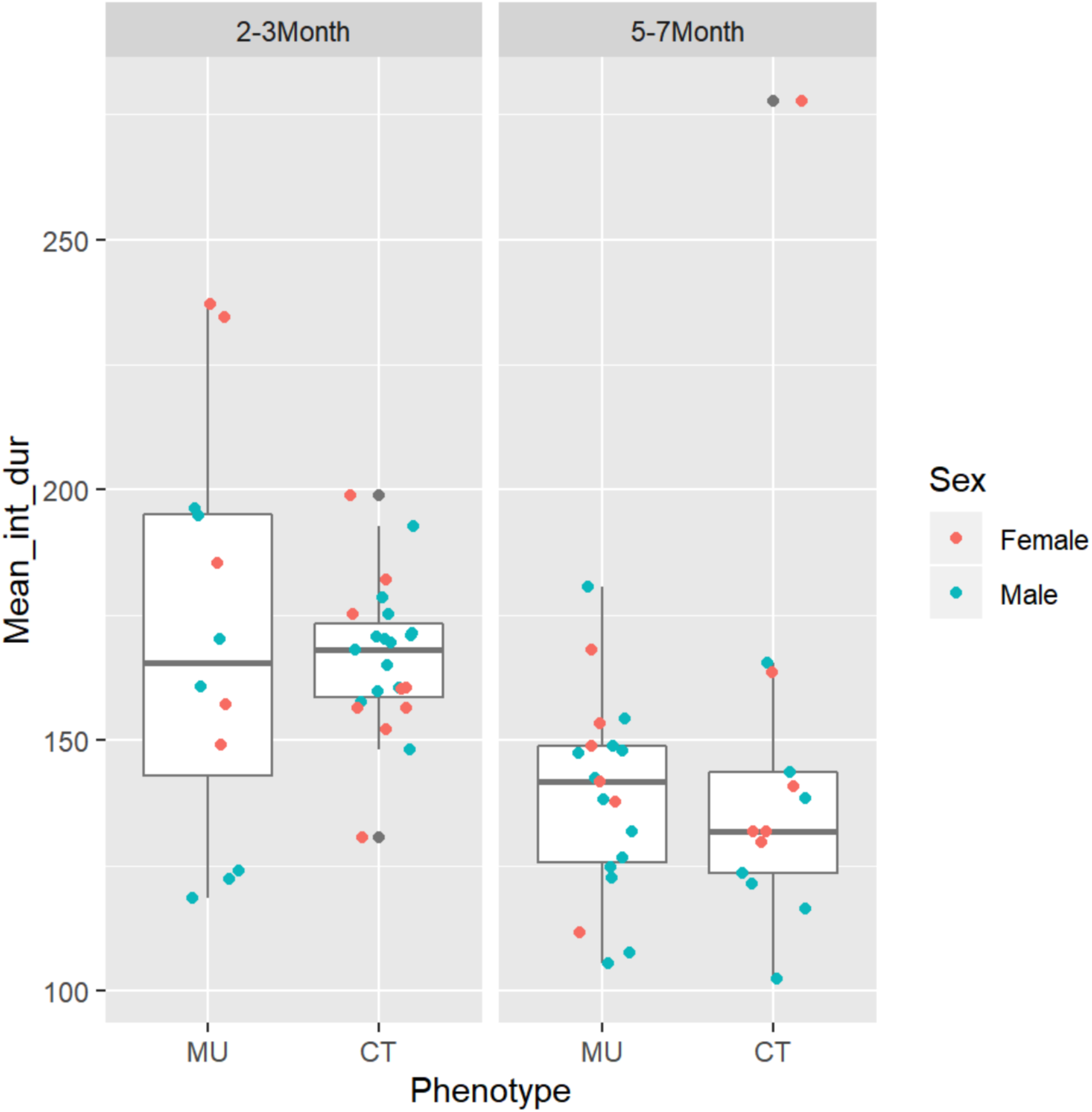
Comparison of mean respiratory interval durations. Mean interval duration is lower in older mice (right panel; p<.001), but does not differ between MU and CT mice within each age-group (left and right box within each panel; p = 0.99, adjusted for age and sex). There seems to be a systematic albeit modest difference by sex (p = 0.048, adjusted for age).

The overall variability of the respiratory rhythm, quantified as the coefficient of variation of the inter-inspiration interval distribution (CV = standard deviation of interval distribution/mean of interval duration) was, like the mean interval duration, dependent on age (p<0.001, with lower values for younger mice), but not on sex (p=0.70). Also in analogy to the mean interval duration above, we could not find a difference between MU mice and their CT littermates with respect to CV (p=0.95, adjusted for age) (Fig. 3).

**Figure 3:**
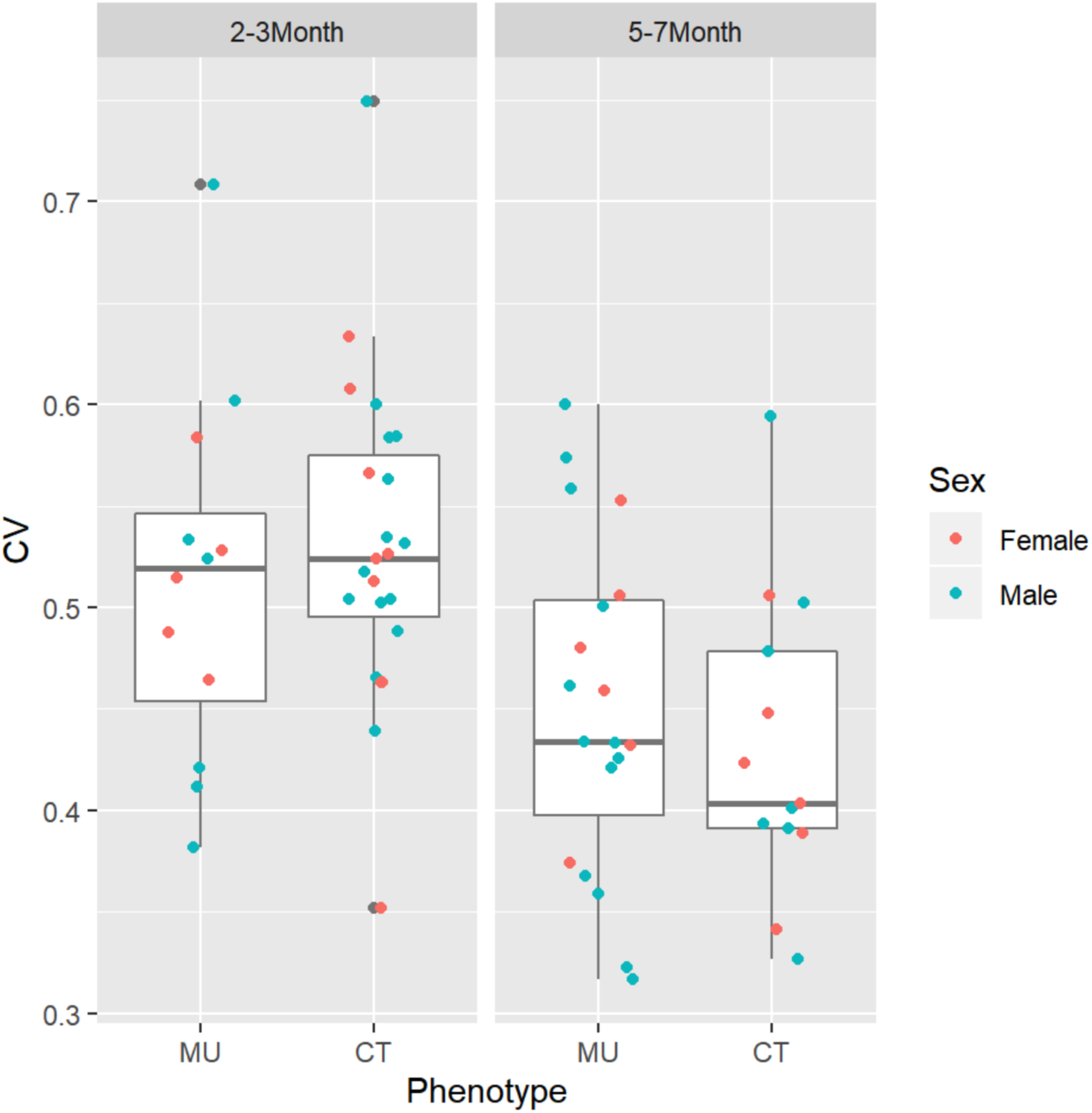
Comparison of the coefficient of variation (CV) of the respiratory interval distribution. The CV is lower in older mice (left vs. right panel; p<.001) but does not differ between MU and CT mice within each age-group (left and right box within each panel; p = 0.95). No systematic differences in the CV were detected by sex (p = 0.70).

The CV measures variability based on the distribution of intervals across the entire observation time. Brief but reoccurring changes in rhythmicity cause no change in the CV, as long as the overall interval distribution remains the same or similar. In 1996, Holt et al. introduced a novel measure of variability, the CV2 (see methods), specifically designed to detect the brief, reoccurring changes in rhythmicity, which they called “intrinsic rhythmicity” (Holt et al., 1996).

Comparison of the CV2 of respiratory behavior in CT and MU mice revealed a lower CV2 in MU mice compared to the CT littermates (p < 0.001), when adjusting for age (p<0.001) and sex (p=0.048) (Fig. 4). While the CV2 increased with age in the control mice (p<0.001), it did not change substantially with age in *L7^Cre^;Vgat^flox/flox^* mice (p=0.16) (Fig. 4). Irrespectively, in the combined data, the interaction effect between age and phenotype was not significant (p = 0.27); something that would be expected if the age-CV2 relationship holds for only one of the two genotypes.

**Figure 4:**
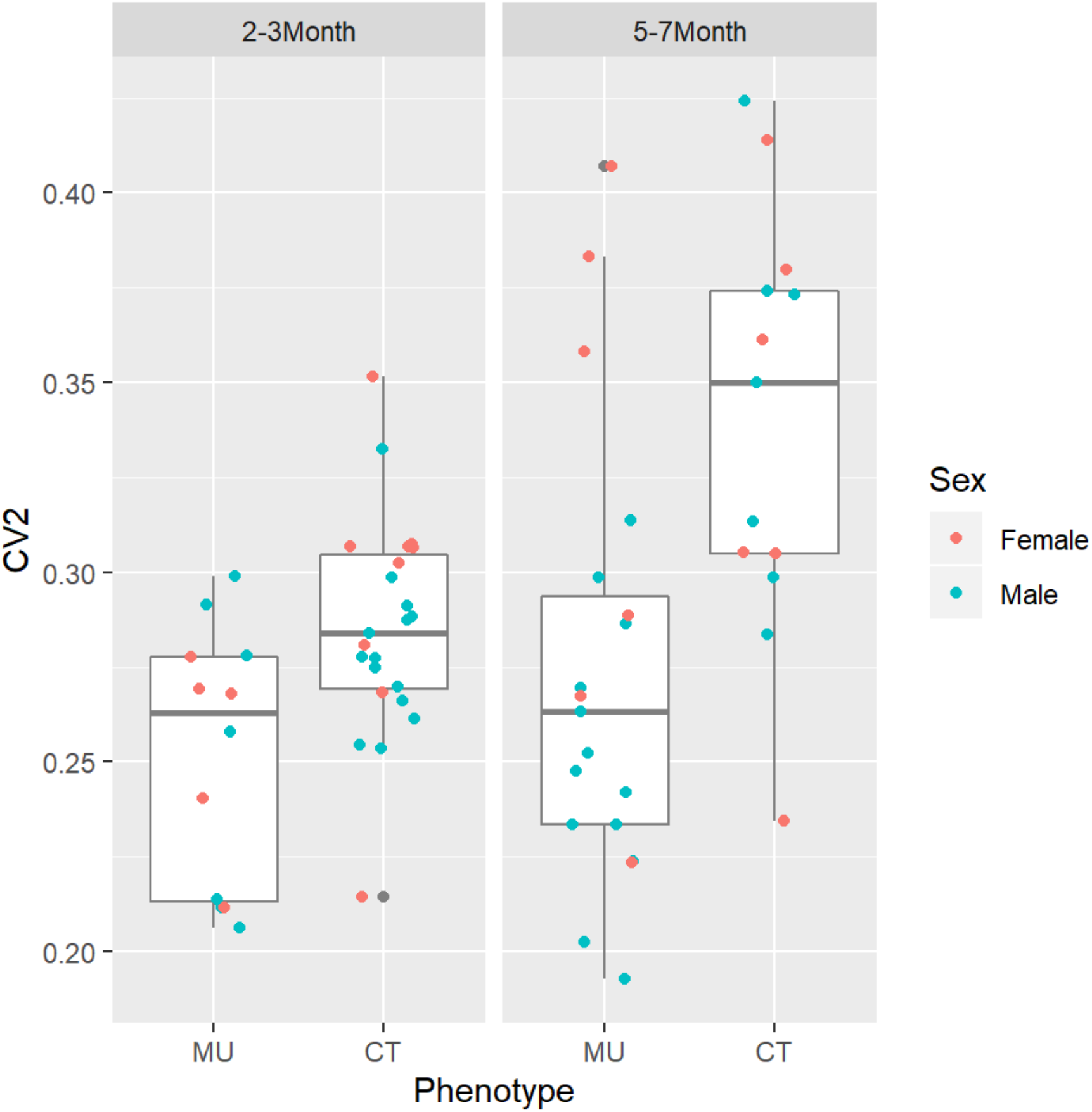
Comparison of the CV2 (intrinsic rhythmicity) of respiratory behavior. The CV2 is lower in the MU mice compared to their CT littermates (left vs. right box in each panel; p<.001) when adjusting for age (p<.001) and sex (p = 0.048). In addition, the CV2 was significantly higher in older compared to younger CT mice (p<.001 in the CT subgroup), but possibly remained unchanged across age in the MU mice (p = 0.16 in the MU subgroup). However, in the combined data, the interaction effect between age and phenotype is not statistically significant (p = 0.19); something that would be expected if the age-CV2 relationship holds for only one of the two phenotypes.

The reduced CV2 in MU mice suggests that the difference in the respiratory sequences in MU and CT mice is based on the temporal modulation of a subset of individual respiratory intervals (see also discussion). In mutant mice, those modulations seem to be less pronounced, resulting in a smaller CV2 value, reflecting reduced intrinsic variability. To address our hypothesis that the difference in CV2 indeed could be explained by modulating the duration of a subset of respiratory intervals, we mimicked in our collected data an extension of the duration of every 10th interval in the respiratory sequence of MU mice by 50%, corresponding to an average of about 75 ms (see methods) (Fig. 5). By so doing, we do render the CV2 values of the respiratory sequence of MU mice more comparable to that of CT mice.

**Figure 5:**
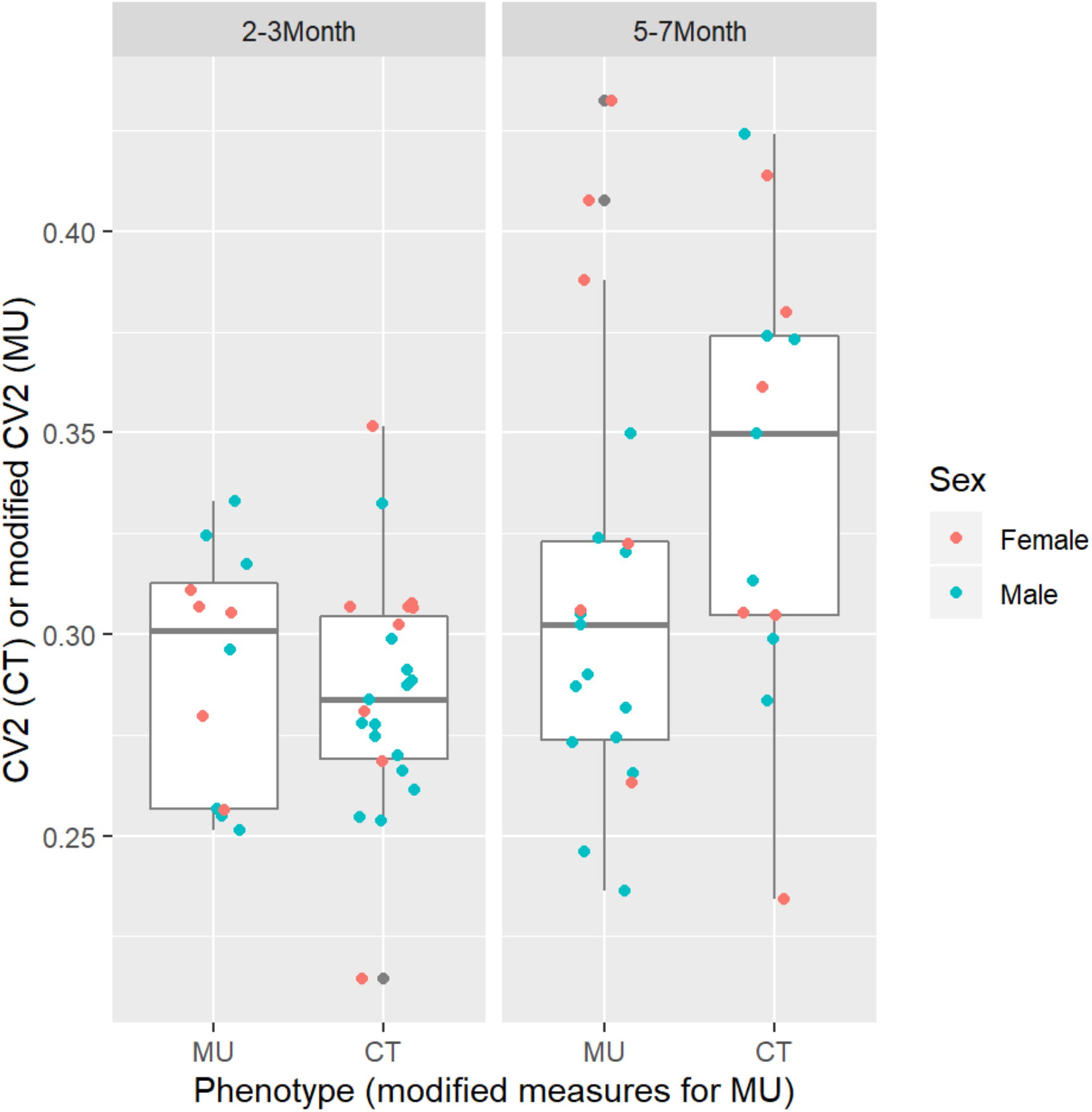
Prolonging the duration of a subset of respiratory intervals to mimic breathing-swallowing coordination in MU mice increased the CV2 to WT values. Using the same analytical approach as in the original respiratory sequence measured in MU mice, one would now conclude that the phenotype is not statistically significant (p = 0.31; age remains significant with p <.001), The respiratory sequences of MU mice was modified *in silico* by extending the duration of every 10^th^ interval by 50%, as described in the methods section.

## DISCUSSION

The cerebellum is known to be part of the neuronal circuitry activated during respiratory challenges, such as hypercapnia or hypoxia (Macey et al., 2005; Parsons et al., 2001). There has been, however, no clear experimental evidence supporting a role of the cerebellum in eupneic breathing (Ebert et al., 1995; Moruzzi, 1940; van der Heijden and Zoghbi, 2018; Xu and Frazier, 2002; Xu et al., 1995). Consistent with previous findings, our results show that loss of cerebellar function does not affect the average respiratory rate or the coefficient of variation of eupneic respiration in mice. Analysis of the CV2, however, showed a significantly reduced variability, i.e. a lower CV2, in MU mice compared to their CT littermates (Fig. 4). The CV2 is uniquely sensitive to brief changes in rhythmicity, which could affect only one or two intervals at a time (Holt et al., 1996). Our findings suggest that loss of cerebellar function is associated with a loss of modulation of select respiratory intervals. A well-known biological purpose for an occasional modulation of respiratory intervals is the coordination of breathing with swallowing movements, which is crucial to protect the airways from aspirating fluid or food (Hardemark Cedborg et al., 2009; Yagi et al., 2017). The typical observation in healthy subjects is an extended pause of the respiratory rhythm during swallowing (swallowing apnea) (Hardemark Cedborg et al., 2009; Nishino, 2012). Inappropriate temporal coordination of swallowing and respiratory movements can lead to dysphagia, a common observation in neurological disorders such as stroke or Parkinson’s disease (Yagi et al., 2017). Based on our observations we propose that the cerebellum is involved in the temporal coordination of breathing and swallowing movements, specifically that the extension of the respiratory pause during swallowing requires an intact cerebellum. We tested this idea *in silico* by artificially prolonging the durations of a subset of respiratory intervals in the respiratory sequences of ataxic mice. Increasing the duration of every 10^th^ respiratory interval by 50% increased the CV2 of the MU sequence to values that no longer differed from CV2 values of CT mice (Fig. 5). At the same time, the changes in average rate and CV caused by this manipulation were insignificant.

Interestingly, it has been reported in humans that breathing-swallowing coordination changes with age (Wang et al., 2015). Older individuals tend to show a reduced frequency of the protective expiration-swallow-expiration pattern, which would increase the risk of aspiration, but accompanying changes, such as longer swallowing apnea, seem to compensate for deficient breathing-swallowing coordination (Wang et al., 2015). While our data suggest an age-related difference in the CV2 of the respiratory sequence, answers to the question of whether a CV2 increase with age is linked to cerebellar function will require a larger sample size size and preferably longitudinal data from the same specimen as they age.

Besides the data we presented here, there are several other lines of evidence supporting a role for the healthy cerebellum in the coordination of respiration with rhythmic orofacial movements. Extensive reciprocal projections connect the cerebellum and the brain stem (Asanuma et al., 1983; Teune et al., 2000; Whiteside and Snider, 1953), providing the necessary anatomical substrate for a cerebellar coordination of brain stem controlled breathing with other orofacial movements. A detailed anatomical and electrophysiological investigation of the medial cerebellar nuclei showed that neurons project broadly to the brain stem, including the area containing the respiratory pattern generating circuits (Lu et al., 2013). Electrophysiological recordings from medial nucleus neurons showed that their spiking activity is correlated with one or two of three different orofacial movements: respiration, fluid licking and mystacial whisker movements (Lu et al., 2013; Xu and Frazier, 2002). In addition, a significant sub population of the medial cerebellar neurons send axon collaterals to more than one brain stem site, a projection pattern that could serve the coordination of brain stem activity at the two sites. Swallowing was not measured in that study, but several lines of evidence support a role for the cerebellum in controlling swallowing. Dysphagia is a common symptom in patients with cerebellar disease (Ramio-Torrentia et al., 2006). The larynx has a neuronal representation in the primate cerebellum (Lam and Ogura, 1952) and functional magnetic resonance imaging showed increased cerebellar activity during swallowing movements (Suzuki et al., 2003).

If the cerebellum operates in the way we propose and does modulate the pause between specific respiratory intervals to allow swallowing movements, we can make specific predictions about neuronal activity patterns in the cerebellum that should be consistent with this operation. Observations in rats have shown that breathing and fluid licking movements have regular rhythms, but swallowing during fluid licking occurs at more variable intervals, every 4^th^ to 14^th^ (Weijnen et al., 1984). Provided the cerebellum coordinates both rhythms, each one should then be represented in the neuronal activity of cerebellar neurons. Neuronal activity in the cerebellum should thus be correlated with the highly regular licking rhythm but also show non-rhythmic activity changes that only occur during licking behavior, presumably linked to swallowing. Such activity patterns were indeed observed in two distinct populations of Purkinje cells in the cerebellum of head fixed mice during fluid licking (Bryant et al., 2010).

In summary, the findings presented here, together with previously published evidence support a proposed role of the cerebellum in the coordination of respiration with swallowing and possibly other rhythmic orofacial movements, such as mystacial vibrissae movements (Bryant et al., 2010; Lu et al., 2013). It is interesting to consider a more general principle of cerebellar rhythm coordination by phase alignment, that could also apply to the coordination of neuronal rhythms downstream of cerebral cortical structures, as suggested by findings of Popa et al. (Popa et al., 2013) and McAfee et al. (McAfee et al., 2019). This proposed coordination of rhythms adds an important new aspect to the widely accepted role of the cerebellum in timing and temporal coordination of sensorimotor activity during behavior (Braitenberg, 1961; Ivry et al., 2002; Mauk and Buonomano, 2004).

## MATERIALS AND METHODS

### Animals

A total of 67 adult mice, 26 females and 41 males, were used in the present study. We compared two groups of mice, one suffering from cerebellar ataxia due to selective loss of Purkinje cell GABAergic synaptic transmission (mutant (MU): *L7^Cre^;Vgat^flox/flox^*, n = 31) and their unaffected littermates (control (CT): *Vgat^flox/flox^*, n = 36) who served as controls (see also Table 1 for details). Mice were either 2-3 or 5-7 months old. During breeding, the day a vaginal plug was detected was considered embryonic day 0.5 and the day of birth postnatal day 0. Genotyping was performed with standard PCR reactions that detect the presence of the *Cre* and *flox* alleles (White et al., 2014). The experimental protocols were approved by the Institutional Animal Care and Use Committee of The University of Tennessee Health Science Center.

### Measurements of respiratory behavior

Respiratory behavior was monitored for 30 min by placing mice in a plethysmograph consisting of a glass container within which mice could move about freely (Fig. 1). The air in the chamber was continuously replaced by a constant flow (1 l/min) of fresh air, directed into the chamber through an opening in the lid with a second opening in the lid serving as an outlet. While the mice were in the chamber, a box was placed over the chamber, blocking direct light.

Inspiration movements cause the air pressure in the plethysmograph chamber to slightly increase. These pressure changes were measured with a pressure transducer (Validyne Engineering, USA). The voltage output of the transducer reflected pressure increase as a decrease in voltage. A decrease in the raw voltage data (example shown in Fig. 1) thus corresponds to inhale movements, and the minima of voltage troughs correspond to the end-of-inhale movements. Voltage data were digitized at 2 kHz using an analogue to digital converter (CED 1401, Cambridge Electronic Design, UK) and stored on computer hard-disk for off-line analysis using signal processing software (Spike 2, Ver. 7, Cambridge Electronic Design, UK). The minima of each voltage trough in the raw data were detected and the times of their occurrences were used as temporal markers of end-of-inspiration. All data analysis was based on the resulting time series of end-of-inspiration markers.

### Statistical analysis

Further analysis of different aspects of the respiratory rhythm, such as mean interval duration, coefficient of variation (CV = standard deviation/mean), and intrinsic variability (CV2) were based on the end-of-inspiration times. The CV2 provides a measure of the similarity of two adjacent inter-inspiration intervals (Equation. 1):

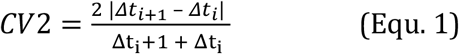

Descriptive boxplots including all animals by phenotype as well as age at time of testing and sex are shown in Fig. 2. Linear models were fitted relating the respective outcome measure to phenotype while adjusting for age and sex. Model fit was verified by residual plots (such as quantile-quantile plots) to verify approximate normality of residuals (not shown). The most important conclusions for CV2 were verified by a bootstrap resampling approach (not shown). Because respiratory function might be affected by age and/or sex, it is important to adjust for these factors when evaluating systematic differences with respect to phenotype. We do this by adding factors for sex and age-group in our regression approach (Kutner et al., 2005). We follow the common approach to exclude a factor if it is not statistically significant (p <= 0.05) because each additional factor reduces the degrees of freedom in the testing for the phenotype-difference, which is our main focus.

### Artificial Modulation of Respiratory Interval Sequences

To mimic a proposed cerebellar dependent extension of respiratory interval duration during swallowing we artificially introduced a predicted cerebellar extension of select respiratory intervals into the respiration sequences measured in MU mice. Based on published findings described in detail below we increased the duration of every 10^th^ respiratory interval by 50%, which translated to an average increase of around 75 ms.

## Acknowledgements

We would like to thank the Neuroscience Institute of the University of Tennessee Health Science Center (UTHSC) for financial support and Micheal Nguyen from the UTHSC Bio-Medical Services for technical support.

## Competing Interests

The authors declare no competing or financial interests.

## Author contribution

D.H.H. and Y.L. conceived and designed the experiments, wrote and edited the manuscript. S.Q. Performed experiments. S.Q. and D.H.H. performed initial data analysis. R.V.S. generated and provided the mutant mouse, wrote and edited the manuscript. B.L.C. and A.P.T. performed data analysis and edited the manuscript. F.T. designed and performed statistical analysis, wrote and edited the manuscript.

## Funding

Funding: D.H.H., Y.L., B.L.C. and A.P.T. are supported by R01MH112143 and R37MH085726. R.V.S. is supported by R01NS089664, R01NS100874, R01MH112143 and U54HD083092.

